# Analysis of the Resuscitation-Availability of Viable-But-Nonculturable Cells of *Vibrio parahaemolyticus* upon Exposure to the Refrigerator Temperature

**DOI:** 10.1101/294751

**Authors:** Jae-Hyun Yoon, Young-Min Bae, Buom-Young Rye, Chang-Sun Choi, Sung-Gwon Moon, Sun-Young Lee

## Abstract

Major pathogenic strains of *Vibrio parahaemolyticus* can enter into the viable-but-nonculturable (VBNC) state when subjected to environmental conditions commonly encountered during food processing. Especially, VBNC cells can be recovered to the culturable state reversibly by removing the causative stress, expressing higher levels of virulence factors. Therefore, the aim of this study was to determine if VBNC *V. parahaemolyticus* strains retain the resuscitation-availability upon eliminating the adverse condition, followed by the enrichment in developed resuscitation-facilitating buffers. Bacterial cells were shown to enter into the VBNC state in artificial sea water (ASW, pH 6) microcosms at 4°C within 70 days. VBNC cells were harvested, inoculated in formulated resuscitation-buffers, and then incubated at 25°C for several days. TSB (pH 8) supplemented with 3% NaCl (TSB_A_) exhibited the higher resuscitation-availability of VBNC cells. It was also shown that TSB_A_ containing 10,000 U/mg/protein catalase, 2% sodium pyruvate, 20 mM MgSO_4_, 5 mM ethylenediaminetetraacetic acid (EDTA), and cell free supernatants extracted from the pure cultures of *V. parahaemolyticus* was more effective in resuscitating VBNC cells of *V. parahaemolyticus*, showing by 7.69-8.91 log_10_ CFU/ml.

**IMPORTANCE:** Generally, higher concentrations (≤40%) of NaCl are used for preserving different sorts of food products from bacterial contaminations. However, it was shown from the present study that strains of *V. parahaemolyticus* were able to persist in maintaining the cellular viability, thereby entering into the VBNC state upon exposure to the refrigerator temperature for 80 days. Hence, the ability of VBNC *V. parahaemolyticus* to re-enter into the culturable state was examined, using various resuscitation buffers that were formulated in this study. VBNC cells re-gained the culturability successfully when transferred onto the resuscitation-buffer D, and then incubated at 25°C for several days. Resuscitation-facilitating agent D is consisting of antioxidizing agents, mineral, an emulsifier, and cell free supernatants from the actively growing cells of *V. parahaemolyticus*. It appeared that such a reversible conversion of VBNC cells to the culturable state would depend on multiple resuscitation-related channels.

## INTRODUCTION

Major food-borne pathogens, including *Vibrio parahaemolyticus*, *Vibrio vulnificus*, *Camphylobacter jejuni*, *Escherichia coli* O157:H7, *Salmonella enterica* serovar Enteritidis, and *Shigella dysenteriae*, are known to become the viable-but-nonculturable (VBNC) state when challenged by various environmental stresses such as low temperatures (≤15°C), starvation, copper, and CO_2_ (1-3). It should be noted that VBNC cells of these pathogens are incapable of producing their own colonies on culture media on which these organisms can grow routinely, thereby escaping from the cultivation-based surveillances and diagnosis tools. Once bacterial cells were induced to the dormant and nonculturable state upon exposure to adverse environmental stresses (nutrient-deprivation and cold temperature) VBNC cells exerted some metabolic activities, including hydrolysis of energy sources, adenosine triphosphate (ATP) synthesis, and maintenance of the membrane integrity, displaying better resistances to environmental conditions commonly encountered during food processing (4-6). Of much importance, it has been well-reported that VBNC *V. parahaemolyticus* can be recovered back to the culturable state by eliminating the causative environmental conditions. Several studies showed that strains of *V. parahaemolyticus* and *V. vulnificus* in such a dormant state were converted to the culturable state on solid agar plates, followed by culturing these long-term-stressed cells in liquid nutrient-rich media at ambient temperatures for several days (4, 7-8). In particular, it was demonstrated that pathogenic bacteria, including *V. parahaemolyticus* and *Shig. dysenteriae*, remained constant in possessing potential virulence factors even after entering into the VBNC state, retaining the serious infectivity to animal cell lines (9-10). Thus, VBNC pathogens should be closely implicated with causing the food-borne disease outbreaks. Until now, many studies have been conducted to determine a way of restoring stressed cells of bacteria from the VBNC state. In a study conducted by Zhao et al. (3), VBNC *E. coli* O157:H7 was transferred into a nutrient-rich culture broth such as tryptic soy broth, and then incubated at 37°C for ≤24 hrs, thereby re-gaining the colony-forming capability. Coutard et al. (11) also showed that VBNC cells of *V. parahaemolyticus* VP5 were resuscitated reversibly when further incubated in artificial sea water (ASW) microcosms at 25°C for several days. In contrast, some VBNC bacteria could not be restored from the VBNC state under controlled favorable conditions where these organisms prefer to grow primarily (12-13). Such a failure to recover VBNC bacteria back to the culturable state did not indicate that the environmental challenges used in these studies deprived bacterial cells of the resuscitation-availability completely. It seemed plausible that these resuscitation approaches would not be effective for recovering the culturability of VBNC cells. Bacteria in such a dormant state will be resuscitated opportunely under a favorable environmental condition for their survivals. Considering that bacterial cells in the VBNC state are capable of evading from conventional cultivation-based techniques the incidence of VBNC pathogens on food products could threaten public health concerns potentially. Until now, a preliminary research establishing an optimal resuscitation method of VBNC cells is still unsubstantial. Therefore, the present study aimed at examining the resuscitation-availability of VBNC *V. parahaemolyticus* using by developed resuscitation-facilitating buffers.

## RESULTS AND DISCUSSION

### Formation of the viable-but-nonculturable cells

It appeared that strains of *V. parahaemolyticus* ATCC 17082, *V. parahaemolyticus* ATCC 33844, and *V. parahaemolyticus* ATCC 27969 were divested of their own culturable capability within 70 days when incubated in ASW microcosms (pH 6) at 4°C, regardless of the excessive amounts of NaCl (Fig. 1). Populations of *V. parahaemolyticus* ATCC 17082 in ASW microcosms containing 0.75%, 5%, 10%, and 30% NaCl declined remarkably below the detection limits (<1.0 log_10_ CFU/ml) as being measured at 4°C for 50, 24, 20, and 12 days, respectively. Especially, such a cold-starvation environment enabled cells of *V. parahaemolyticus* ATCC 33844 and *V. parahaemolyticus* ATCC 27969 to be converted into the nonculturable state in ASW microcosms amended with 30% NaCl for ≤24 days. In addition, these organisms became nonculturable when incubated in ASW microcosms added with less than 10% NaCl at 4°C for 70 days. Clearly, there were minor modifications in the duration of cold-starvation periods required for these pathogens to lose the 100% culturability. Nevertheless, it seemed likely clear that *V. parahaemolyticus* were converted to the nonculturable state more rapidly with increasing NaCl concentrations. Furthermore, it was shown that viable numbers of *V. parahaemolyticus* ATCC 17082, *V. parahaemolyticus* ATCC 33844, and *V. parahaemolyticus* ATCC 27969 ranged from 4.3 to 6.5 log_10_ CFU per a slide after incubated at 4°C for 80 days with the fluorescence microscopic assay. Apart from the culturable populations of these bacteria, strains of *V. parahaemolyticus* persisted in surviving under the cold-starvation condition for at least 80 days. In order to determine whether the nonculturable cells were truly dead or still alive, it is inevitable to evaluate the degree to which these organisms were sincerely damaged. Then, the utilization of fluorescent probes such as SYTO9 and propidium iodide (PI) can reflect the levels of cellular integrity quantitatively. In general, SYTO9 penetrates bacteria with the intact cell membrane, interacts with the cell nucleic acid, and then displays green colors for the live cells with the fluorescence microscopy. Propidium iodide can penetrate damaged membranes, labeling only the dead cells as red-coloured fluorescence. Hence, staining bacterial cells with SYTO9 combined with PI can distinguish between live and dead cells effectively. Herein, these results indicated that cells of *V. parahaemolyticus* were inducted into the VBNC state, still maintaining the cellular integrity of bacterial membranes even after a long term of cold-starvation

**FIG. 1.**
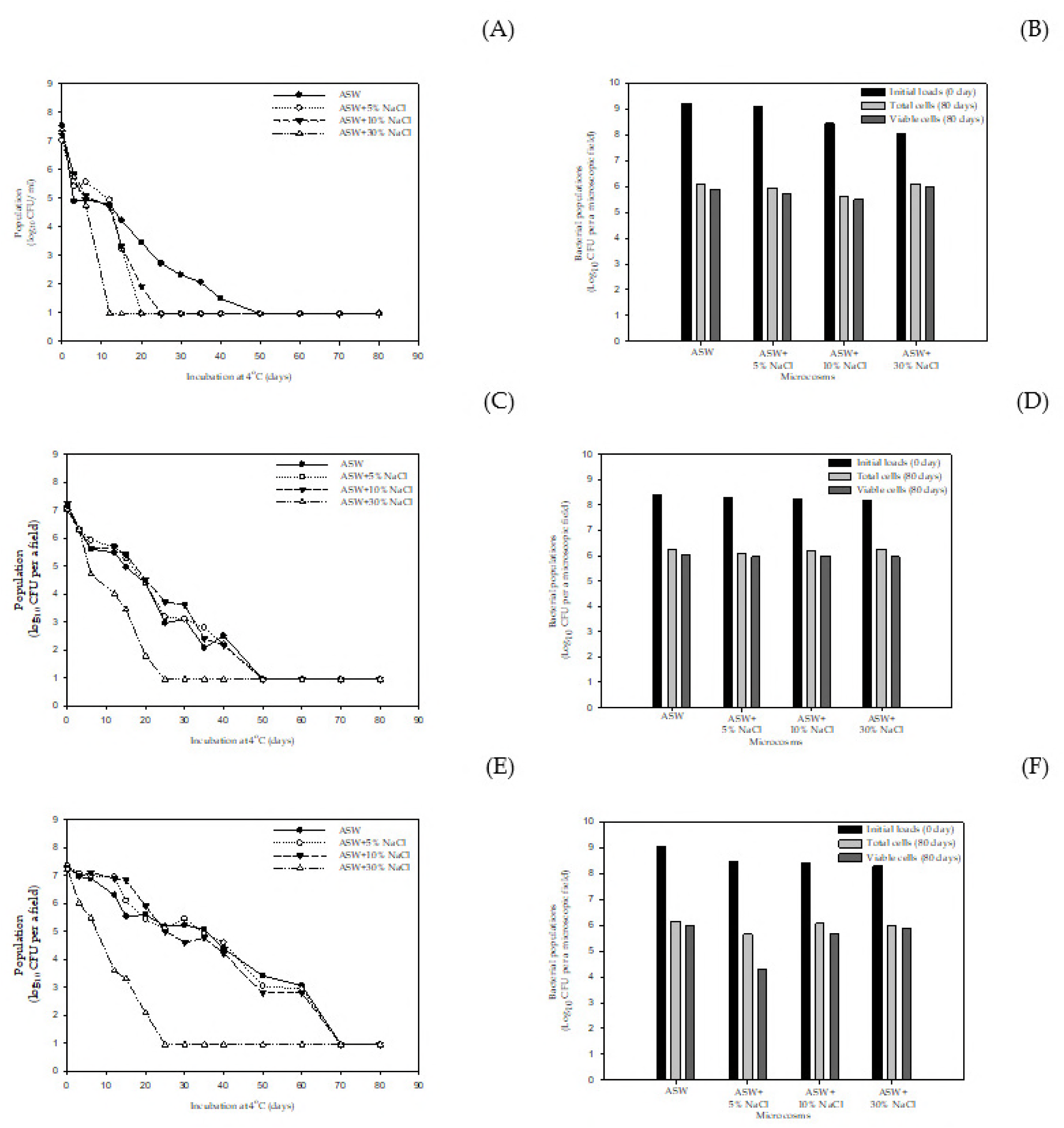
Loss of the culturability (A, C, and E) and the viability (B, D, and F) of *V. parahaemolyticus* ATCC 17082 (A-B), *V. parahaemolyticus* ATCC 33844 (C-D), and *V. parahaemolyticus* ATCC 27969 (E-F) incubated in ASW (pH 6) microcosms supplemented with varying concentrations of NaCl at 4°C for 80 days.

**Fig. 2.**
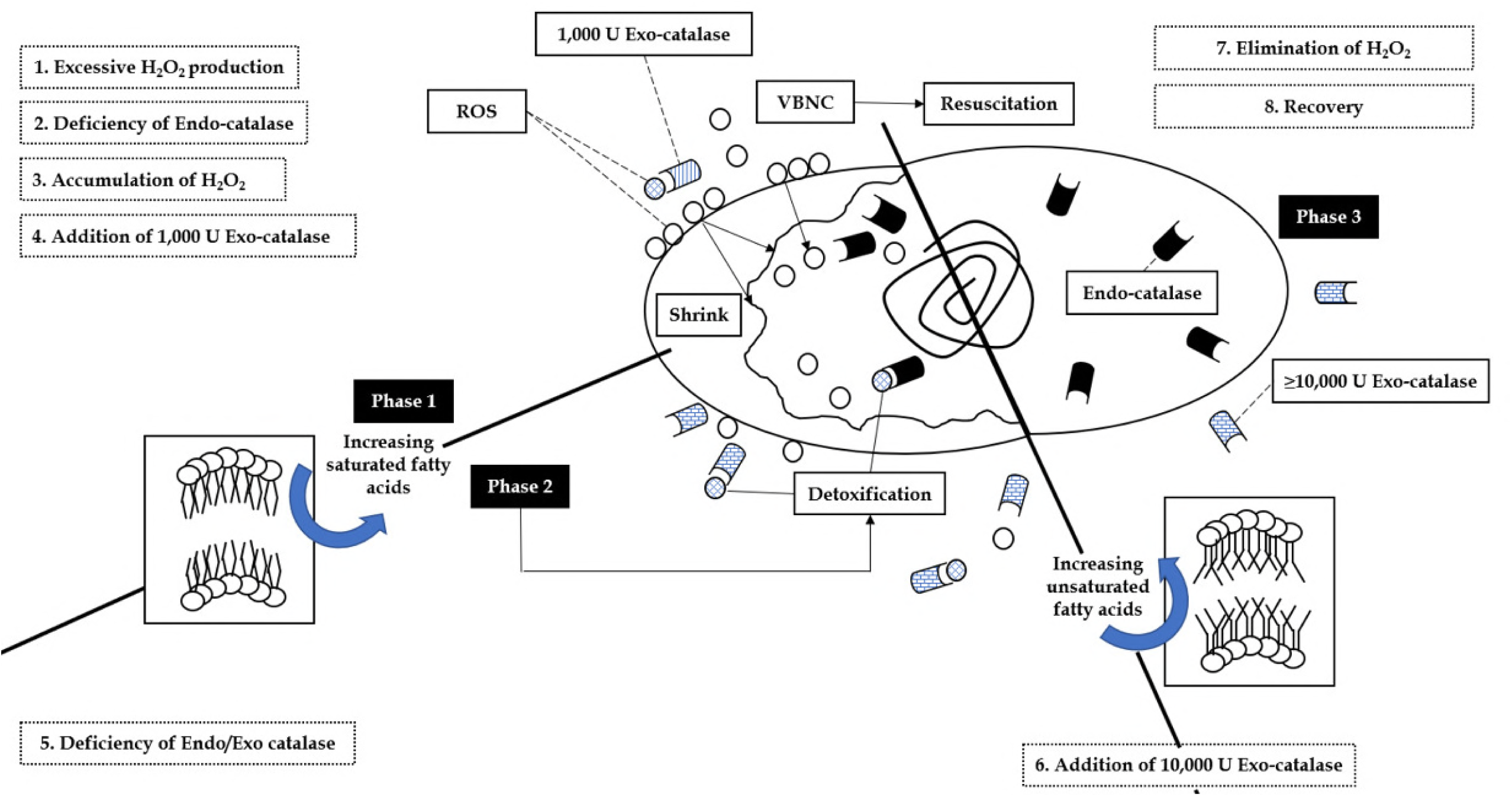
A hypothesis diagram for elucidating the successful recovery of VBNC cells back to the re-culturable state by the addition of a high degree of (>10,000 U/mg/protein) of catalase proteins (Exo-catalase) in the resuscitation buffer.

As well-known previously, *V. parahaemolyticus* strains are very susceptible to various environmental conditions. Probably, it would be attributable to the low levels of detectability of these bacteria on food products. It was shown that bacterial cells of *V. parahaemolyticus* were derived from the colony-forming capability on culture media after the entry into the VBNC state (19). In general, pathogenic bacteria such as *V. parahaemolyticus*, *V. vulnificus*, and *V. cholerae* can enter into the VBNC state at low temperatures of less than ≤10°C within a wide range of incubation periods. As shown in Table 1, it was demonstrated that these pathogens can be converted into the VBNC state by various environmental conditions. Baffone et al. (14) reported that a cold-starvation challenge enabled cells of *V. parahaemolyticus* and *V. vulnificus* to enter into the VBNC state within 30 days. Similarly, strains of *V. parahaemolyticus* and *V. vulnificus* were converted into the VBNC state when incubated in ASW microcosms at 4°C within 35 days (11, 18). Moreover, these bacteria became viable-but-nonculturable successfully in various microcosms such as ASW, deionized water (DW), natural sea water, and a mixture of ASW and a Luria-Bertani culture medium. Interestingly, it appeared that strains of *V. parahaemolyticus*, *V. vulnificus*, and *V. cholerae* required very different incubation-periods to enter into the VBNC state under the conditions that are almost the same as proposed in previous studies. The duration of cold-starvation stress ranged from 4 days to a maximum of several months to produce VBNC cells. Although the underlying mechanisms governing the entrance of *V. parahaemolyticus* into such a dormant state has not been understood yet, the formation of VBNC cells would proceed by a multiple mode of actions and complex interactions either directly or indirectly. Clearly, the formation of VBNC cells should be recognized as one of the adaptation-surviving strategies in response to adverse environments.

**TABLE 1.** Effects of environmental conditions on the entry of *V. parahaemolyticus*, *V. vulnificus*, and *V. cholerae* into the viable-but-nonculturable

| Microorganism | VBNC-inducing conditions |  |  |  |  | Resuscitation | Reference |
| --- | --- | --- | --- | --- | --- | --- | --- |
|  | Major stress | Microcosm | Temperature | Period | Media<sup>2</sup> |  |  |
| *Vibrio parahaemolyticus* | Temperature/Starvation | ASW | 5°C | 30 | BHI | ND | (14) |
| *Vibrio parahaemolyticus* | Temperature/Starvation | ASW | 4°C | 50 | TSA | ○ | (8) |
| *Vibrio parahaemolyticus* | Temperature/Starvation | ASW | 4°C | 12 | LB | —<sup>3</sup> | (15) |
| *Vibrio parahaemolyticus* | Temperature/Starvation | ASW | 5°C | 4 | HIA | ○ | (25) |
| *Vibrio parahaemolyticus* | Temperature/Starvation | MMS | 3.5°C | 50-70 | TCBS | - | (26) |
| *Vibrio parahaemolyticus* | Temperature/Starvation | ASW | 10°C | <30 | HIA | ○ | (11) |
| *Vibrio parahaemolyticus* | Temperature/Starvation | MMS | 4°C | 35 | TSA | ○ | (9) |
| *Vibrio vulnificus* | Temperature/Starvation | ASW | 4°C | 14 | Tn | ○ | (21) |
| *Vibrio vulnificus* | Temperature/Starvation | ASW | 5°C | 4 | HIA | ○ | (16) |
| *Vibrio vulnificus* | Temperature/Starvation | ASW | 5°C | 10 | HIA | ○ | (17) |
| *Vibrio vulnificus* | Temperature/Starvation | ASW | 5°C | <30 | LBA | ND | (18) |
| *Vibrio cholerae* | Temperature/Starvation | ND | 4°C | 70 | ND | ND | (20) |
| *Vibrio cholerae* | Temperature/Starvation | ASW | 4°C | 20-30 | TSA | ND | (22) |
| *Vibrio cholerae* | Temperature | ASW+LB | 4°C | 30-40 | TSA | ND | (22) |
| *Vibrio cholerae* | Anaerobic atmosphere | ASW | 4°C | 40 | TSA | ND | (22) |
| *Vibrio cholerae* | Starvation | Sea water | ND | ≤125 | BHI | ○ | (23) |

<sup>1</sup> BHI, brain heart infusion agar; TSA, tryptic soy agar; LBA, Luria-Bertani agar; HIA, heart infusion agar; NA, nutrient agar; 2216E, marine agar; TCBS, thiosulphate-citrate-bile salt-sucrose agar.
<sup>2</sup> ND, not determined.

### Evaluation of the resuscitation-availability of VBNC cells

After forming VBNC cells of *V. parahaemolyticus* strains, these cells were transferred to various nutrient-rich culture fluids, including ASW, TSB, BHI, and APW, and then further incubated to examine if VBNC cells would retain the resuscitation-availability on these media at an ambient temperature (Table 2-5). It was shown that VBNC cells failed to re-gain the culturability when re-suspended in a formal solution of ASW for several days. However, once resuscitated in nutrient-rich media such as TSB and APW, VBNC cells of *V. parahaemolyticus* ATCC 17082 were turned back to the culturable state, showing by 3.45– 8.00 log_10_ CFU/ml as being enumerated on a nonselective medium (TSA). Strains of *V. parahaemolyticus* ATCC 33844 and *V. parahaemolyticus* ATCC 27969 also resuscitated positively when bacterial cells in the VBNC state were incubated in TSB and APW, except for these organisms that had been induced into the VBNC state in ASW microcosms added with 30% NaCl at 4°C for 80 days. Resuscitation-provoking efficiencies of these buffers such as TSB and APW were in the levels of ≥6.0 log_10_ CFU/ml, whereas the selection of BHI was less effective for the resuscitation of VBNC *V. parahaemolyticus*. Therefore, TSB as a resuscitation-buffer facilitated the recovery of VBNC cells of these pathogens, indicating that certain levels of a minimum nutritional base should be required for VBNC *V. parahaemolyticus* to be recovered to the culturable state. As far as it is controversial to determine the effects of CFS extracted from major food-borne pathogens and a mixture of antioxidizing agents, CSP, on the resuscitation of VBNC *V. parahaemolyticus* VBNC cells resuscitated in CFS-VP showed the colony-forming capability, ranging from 7.50 to 8.38 log_10_ CFU/ml. The use of CFS-VV was also attributable to the moderate recovery of VBNC cells, but showed lower levels of the resuscitation-availability than that of CFS-VP. VBNC cells of *V. parahaemolyticus* ATCC 33844, which were challenged by cold-starvation in ASW microcosms added with 5% NaCl for 80 days previously, were not awakened to the culturable state, followed by the resuscitation process to CFS-EC, CFS-ST, and CFS-SA, respectively. These results were in an accordance with a study conducted by Pinto et al. (24). Ayrapetyan et al. (27) also showed that VBNC cells of *V. vulnificus* were able to be awakened from such a dormant state when resuscitated on culture media supplemented with the CFS extracted from the pure cultures of *V. vulnficus*. Preliminarily, it was revealed out that autoinducer-2 (AI-2) could be strongly involved in the resuscitation of VBNC *V. vulnificus*, whereas filtered CFSs from AI-2 mutant strains of *V. vulnificus* failed to restore VBNC cells. These results implied that interspecific quorum sensing modules would play a key role as an important regulator in switching on the resuscitation-availability of VBNC bacteria. In our preliminary studies, several intrinsic parameters such as pH and NaCl% in TSB were adjusted to establish an optimal condition in an attempt to initiate the resuscitation of VBNC *V. parahaemolyticus* and VBNC *V. vulnificus* (*data not shown*). Consequently, TSB (pH 8) supplemented with 3% NaCl (TSB_A_) exhibited the higher resuscitation-availability of VBNC cells of these pathogens. Unexpectedly, APW (pH 7-8) solutions amended with 1%-3% NaCl were not effective in awakening the restoration of VBNC cells. Thus, 100-days-stressed cells of *V. parahaemolyticus* in ASW microcosms at 4°C were transferred to either TSB_A_+CFS or TSB_A_+CSP, showing that VBNC cells of *V. parahaemolyticus* ATCC 17082 were not converted to the culturable state while strains of *V. parahaemolyticus* ATCC 33844 and *V. parahaemolyticus* ATCC 27969 in ASW microcosms added with ≤10% NaCl re-gained the culturability, corresponding to 7.69-8.91 log_10_ CFU/g (Table 2). Especially, TSB_A_+CFS-VP was more effective in resuscitating VBNC *V. parahaemolyticus* than TSB_A_ combined with CFS-VV, CFS-EC, CFS-ST or CFS-SA. Based on these results, buffers A-F were prepared to determine the optimal resuscitation-buffer (Table 4). Above all things, 250-days-stressed cells of *V. parahaemolyticus* ATCC 33844 in ASW microcosms added with 30% NaCl were not awakened from such a dormant state with resuscitation-buffers A-F. However, resuscitation-buffer D facilitated VBNC cells of *V. parahaemolyticus* ATCC 17082 and *V. parahaemolyticus* ATCC 27969 to re-gain the colony-forming capability effectively. It was shown that a mixture of CFS-VP, CSP, MgSO_4_, and EDTA was highly effective in resuscitating VBNC cells.

**TABLE 2.** Resuscitation (log_10_ CFU/ml) of *V. parahaemolyticus* from the VBNC state with the temperature upshift at 25°C for 3 days

| Pathogen | ATCC | Microcosm | Cold-starvation period (days)<sup>1</sup> | Resuscitation-buffer |  |
| --- | --- | --- | --- | --- | --- |
|  |  |  |  | ASW | TSB |
| *V. parahaemolyticus* | 17082 | ASW | 50 | -<sup>2</sup> | 7.44 |
|  | 17082 | ASW+5% NaCl | 25 | - | - |
|  | 17082 | ASW+10% NaCl | 21 | - | - |
|  | 17082 | ASW+30% NaCl | 12 | - | - |
| *V. parahaemolyticus* | 33844 | ASW | 50 | - | 7.06 |
|  | 33844 | ASW+5% NaCl | 50 | - | 5.70 |
|  | 33844 | ASW+10% NaCl | 50 | - | - |
|  | 33844 | ASW+30% NaCl | 25 | - | - |
| *V. parahaemolyticus* | 27969 | ASW | 72 | - | 7.44 |
|  | 27969 | ASW+5% NaCl | 72 | - | - |
|  | 27969 | ASW+10% NaCl | 72 | - | - |
|  | 27969 | ASW+30% NaCl | 25 | - | - |

<sup>1</sup>*V. parahaemolyticus* and *V. vulnificus* were incubated in ASW (pH 6) microcosms supplemented with varying levels of NaCl at 4°C until they became the VBNC state. In particular, these pathogens in the VBNC state for the first day were transferred onto these resuscitation-buffers such as ASW and TSB, respectively, and then they were incubated at 25°C for 72 hrs.
<sup>2</sup> -, No growth.

**TABLE 3.**
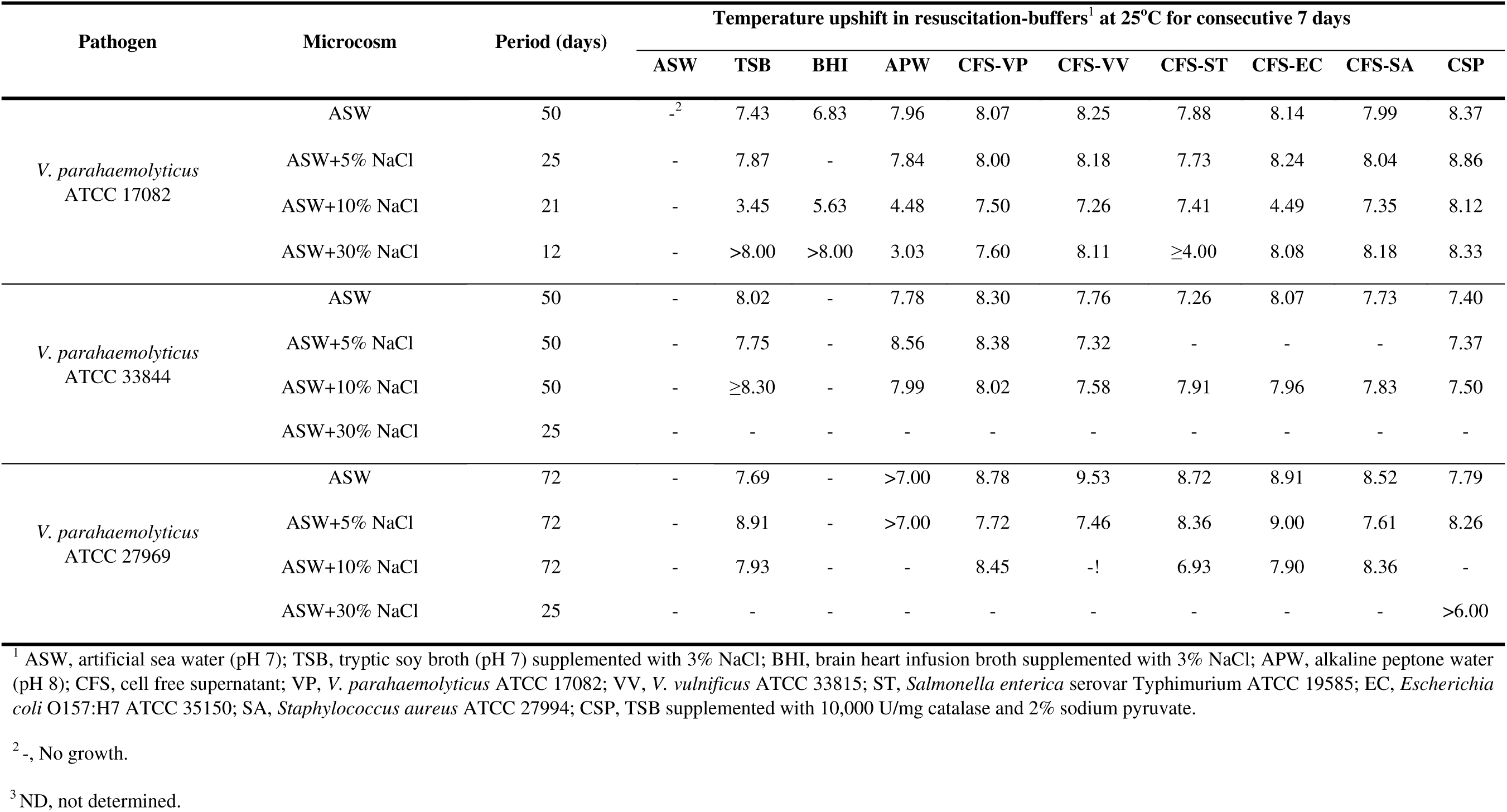
Evaluation of the ability (log_10_ CFU/ml) of *V. parahaemolyticus* pre-incubated in ASW microcosms (pH 6) at 4°C for 60 days to be recovered from the VBNC state

**TABLE 4.** Effects of NaCl contents (1-3%) combined with alkaline pH levels (pH 8-9) on the resuscitation of *V. parahaemolyticus* in the VBNC state at 4°C for 80 days

| Pathogen | Microcosm | Temperature upshift in the following resuscitation-buffers<sup>1</sup> at 25°C for consecutive 7 days |  |  |  |  |  |  |  |
| --- | --- | --- | --- | --- | --- | --- | --- | --- | --- |
|  |  | TSB | TSB 1 | TSB 2 | TSB 3 | APW | APW 1 | APW 2 | APW 3 |
| *V. parahaemolyticus*<br>ATCC 17082 | ASW | ○ | ○ | ○ | - | - | - | - | - |
|  | ASW+5% NaCl | -<sup>2</sup> | - | - | - | - | - | - | - |
|  | ASW+10% NaCl | - | - | - | - | - | - | - | - |
|  | ASW+30% NaCl | - | - | - | - | - | - | - | - |
| *V. parahaemolyticus*<br>ATCC 33844 | ASW | ○ | ○ | ○ | ○ | - | - | ○ | - |
|  | ASW+5% NaCl | ○ | ○ | ○ | ○ | - | - | ○ | - |
|  | ASW+10% NaCl | - | ○ | ○ | ○ | - | - | - | - |
|  | ASW+30% NaCl | - | - | - | - | - | - | - | - |
| *V. parahaemolyticus*<br>ATCC 27969 | ASW | ○ | ○ | ○ | ○ | ○ | ○ | ○ | ○ |
|  | ASW+5% NaCl | ○ | ○ | ○ | - | ○ | ○ | ○ | ○ |
|  | ASW+10% NaCl | ○ | - | ○ | - | ○ | ○ | ○ | ○ |
|  | ASW+30% NaCl | - | - | - | - | - | - | - | - |

<sup>1</sup> TSB, tryptic soy broth (pH 7) supplemented with 3% NaCl; TSB 1, TSB (pH8) supplemented with 3% NaCl; TSB 2, TSB (pH 7) supplemented with 1% NaCl; TSB 3, TSB (pH 8) supplemented with 1% NaCl; APW, Alkaline Peptone Water (pH 8) supplemented with 3% NaCl; APW 1, APW (pH 8) supplemented with 3% NaCl; APW 2, APW (pH 7) supplemented with 1% NaCl; APW 3, APW (pH 8) supplemented with 1% NaCl.
<sup>2</sup> -, No growth.

To data, it was believed that the intracellular accumulation of reactive oxygen species (ROS) has a significant influence on the formation of VBNC cells of organisms. Accordingly, the incidence of ROS-detoxifying proteins in bacterial cells could be closely associated with the resuscitation-availability, probably preventing the bacteria from entering into the VBNC state. In our preliminary studies, it was shown that there were no significant differences in the amounts of antioxidizing proteins such as catalase and glutathione-S-transferase between the bacterial cells of *V. parahaemolyticus* in the stationary-phase and in the VBNC state (*data not shown*). Once VBNC cells of *V. parahaemolyticus* in ASW microcosms at 4°C for 90 days were harvested by centrifugation at 1,3000 X g for 3 min, washed twice, and then re-suspended in 5 ml of TSA (pH 7) either containing 1,000 U/mg/protein or 10,000 U/mg/protein none of bacterial cells were resuscitated in TSB added with 1,000 U catalase, while these cells re-suspended in TSB+10,000 U catalase were converted back to the culturable state, showing by ≥7.0 log_10_ CFU/ml on media (*data not shown*). It seemed plausible that if accumulated amounts of ROS exceed over an acceptable coverage of ROS-detoxifying enzyme’s activity bacteria begin to proceed towards the VBNC stage (Fig. 1). It was suggested that ROS accumulations would play an important role for understanding related mechanisms governing the entrance of *V. parahaemolyticus* into the VBNC state. After VBNC cells of *Ralstonia solanacearum* were incubated in DW amended with 1,000 U/mg catalase at 30°C for 3 days, this bacterium re-gained the culturability on media, showing by >8.0 log_10_ CFU/ml (28). Mizunoe et al. (15) reported that VBNC cells of *V. parahaemolyticus* was plating-counted on TSA (pH 7) added with 1,000 U/mg protein catalase, thereby resulting in a mild restoration of the resuscitation-availability. Interestingly, Abe et al. (29) revealed out that GST activities of *V. vulnificus* remained constant during cold-starvation for 24 hrs, showing by 3.07-3.83 ưM/mg/protein uniformly. Under the same condition, nonculturable suppression mutant strains of *V. vulnificus* persisted in showing the colony-forming capability in the levels of ≥5.0 log_10_ CFU/ml, and then exerted enhanced GST activities more than 10 times as massive as the pure cultures. In a study conducted by Santander et al. (30), mutant strains of *Erwini amylovora* deleting katAG^−^ entered into the VBNC state in ASW microcosms at 4°C more rapidly than did the wild cells. Each of katA and katG is responsible for producing a monofunction catalase and a bifunctional peroxidase, respectively. Therefore, these publications suggested that the activities of ROS-scavenging agents in bacterial cells could be strongly associated with the loss of culturability. These results, along with our findings, would represent the hypothesis that bacterial cells detoxify intracellularly generated ROS materials by synthesizing catalase/GST-like enzymes at the beginning stage of cold-starvation (Fig. 1). After at least several weeks, bacteria would not synthesize enough amounts of the antioxidizing enzymes to hydrolyze the accumulated ROS, eventually resulting in the loss of culturability. Once bacterial cells became the nonculturable state, VBNC cells were further transferred into nutrient-rich media added with different concentrations (ex. 1,000 or 10,000 U/mg/protein) of catalase, and then incubated at ambient temperatures for several days the resuscitation-buffer reinforced with 10,000 U/mg/protein catalase would provide VBNC cells with a sufficient quantity of the ROS-detoxifying agent successfully to neutralize intracellular ROS materials, allowing bacterial cells to be recovered from the culturable state again despite that VBNC bacteria resuscitated in liquid culture broth containing 1,000 U/mg/protein catalase were incapable of gaining the re-culturability as the existing accumulated amounts of ROS would exceed over those of added Exo-catalase proteins.

## MATERIALS AND METHODS

### Preparation of bacterial inoculums

Strains of *V. parahaemolyticus* ATCC 17082, *V. parahaemolyticus* ATCC 27969, and *V. parahaemolyticus* ATCC 33844 were purchased from the Korean Collection for Type Cultures (KCTC, Daejon, Korea). Bacterial stocks these cells were maintained at −75°C and further activated in tryptic soy broth (Difco, Detroit, MI, USA) supplemented with 3% NaCl (TSB) at 37°C for 24 hrs before use. Stationary phase cells of *V. parahaemolyticus* were harvested by centrifugation at 10,000 X g for 3 min, washed with artificial sea water (ASW, Sigma-Aldrich, St. Louis, MO, USA), and then final pellets of these organisms were re-suspended in 1 ml of ASW solutions (pH 6), corresponding to the bacterial population of approximately 10^8-9^ CFU/ml. To adjust the pH level in ASW microcosms, filtered 1 N NaOH solution (Kanto chemical, Tokyo, Japan) was used.

ASW solutions (Sigma-Aldrich, St. Louis, MO, USA) were prepared according to the manufacturer’s instruction. Formal ASW microcosms (pH 7.2-7.8) contained 19,290 mg of Cl, 10,780 mg of Na, 2,660 mg of SO_4_, 420 mg of K, 400 mg of Ca, 200 mg of CO_3_, 8.8 mg of Sr, 5.6 mg of B, 56 mg of Br, 0.24 mg of I, 0.3 mg of Li, 1.0 mg of F, and 1,320 mg of Mg per 1 liter of sterile distilled water. These microcosms were autoclaved at 125°C for 20 min before use. Then, NaCl concentrations of these microcosms were adjusted to 0.75%, 5%, 10%, and 30% (m/v), respectively. Each of these microcosms was adjusted to pH 6.0–6.2, using a membrane-filtered 1N NaOH solution (Kanto chemical, Tokyo, Japan), facilitating the induction of *V. parahaemolyticus* into the VBNC state. Then, bacterial cells were inoculated in 100 ml of ASW (pH 6) microcosms added with 0.75%, 5%, 10%, and 30% NaCl, respectively. Bacterial suspensions were kept at 4°C until the culturable numbers of *V. parahaemolyticus* arrived below the detectable limits (<1.0 log_10_ CFU/ml). ASW microcosms were withdrawn from the incubator at regular time-intervals to enumerate the bacterial population either directly by the cultivation-based method or indirectly by measuring the viable cell numbers of *V. parahaemolyticus*.

### Enumeration of the bacterial population

Cells of *V. parahaemolyticus* ATCC 17082, *V. parahaemolyticus* ATCC 33844, and *V. parahaemolyticus* ATCC 27969 were plating-counted on typtic soy agar (Difco) supplemented with 3% NaCl (TSA). Decimal dilutions (10^−1^) of the bacterial cell were prepared in alkaline peptone water (APW, Difco) consisting 10 g of peptone and 10 g of NaCl per 1 liter of DW. Then, 100 µl of these aliquots was spread on TSA. Each of these plates were incubated at 37°C for 24 hrs and colonies developed on media were further enumerated.

### Fluorescence dye staining and microscopic assay

Numbers of total and viable cells of *V. parahaemolyticus* were determined as being measured with the Live/Dead^®^ BacLight™ Bacterial Viability Kit (Invitrogen, Mount Waverley, Victoria, Australia) combining two nucleic acid stains, SYTO9 and propodium iodide. It has been well-documented that SYTO9 has a high affinity for deoxyribonucleic acid (DNA) and chromosome of bacterial cells, labelling all of the bacteria with intact and compromised membranes, whereas propodium iodide penetrates selectively the bacterial cell with damaged membranes. Briefly, equal volumes (1:1) of SYTO9 and propodium iodide were combined and 3 µl of this mixture was added to each 1 ml of the bacterial cell. After a short period (ca. 15 min) of incubation at an ambient temperature in the dark, 5-8 µl of this aliquot was attached on a glass slide and a coverslip was placed on this specimen carefully. Then, bacterial images were demonstrated using an electron-fluorescent microscope (TE 2000-U, Nikon, Japan).

### Optimization of the resuscitation-facilitating buffers

In the present study, various resuscitation-facilitating strategies were employed to examine the resuscitation-availability of VBNC cells to the culturable state on media. (Ι) VBNC cells of *V. parahaemolyticus* were centrifugated at 13,000 X g for 3 min, washed twice, and then re-suspended in 5 ml of ASW (pH 7) solutions containing 0.75% NaCl. Immediately, these cells were incubated at 25°C for up to 7 days. (ΙΙ) Several nutrient-rich media, including APW, TSB, and brain heart infusion (BHI, Difco) broth, were prepared according to instructions provided by suppliers. These media were added with excessive amounts of NaCl, corresponding to either 1% or 3% NaCl, and pH levels of these media were adjusted to either pH 7 or pH 8, using a membrane-filtered 1N NaOH solution. VBNC cells were harvested by centrifugation at 13,000 X g for 3 min, washed twice, re-suspended in 5 ml of these culture media, respectively, and then further were incubated at 25°C for up to 7 days. (ΙΙΙ) As shown in Table 4, each of supplementations, including 10,000 U/mg protein catalase (Sigma), 2% sodium pyruvate (Sigma), 20 mM MgSO_4_ (Sigma), 5 mM ethylenediaminetetraacetic acid (EDTA, Sigma), and cell free supernatant (CFS), were individually added to TSB (pH 8) containing 3% NaCl to alter various resuscitation-facilitating buffers. When it comes to CFS, these fluids were extracted from the wild cells of *V. parahaemolyticus* ATCC 17082, *V. vulnificus* ATCC 27562, *Escherichia coli* O157:H7, *Salmonella enterica* serovar Typhimurium, and *Staphyloccus aureus*, respectively. Briefly, each of these pathogens grown in TSB at 37°C for 24 hrs were harvested by centrifugation at 13,000 X g for 3 min to collect bacterial pellets. These supernatants were separately collected, filtered through a 0.2-ưm-size a polycarbonate membrane (ADVANTEC, Tokyo, Japan), and then added in resuscitation-buffers D-F at a ratio of 10% (v/v). It was further confirmed that all the CFSs did not have an influence on the controlled intrinsic pH level in TSB. VBNC cells were centrifugated at 13,000 X g for 3 min, washed twice, and then re-suspended in 5 ml of resuscitation-buffers A-F, respectively. At the end, bacterial cells were incubated at 25°C for up to 7 days.

**TABLE 5.** Assessment of the ability (log_10_ CFU/g) of *V. parahaemolyticus* pre-incubated in ASW microcosms (pH 6) at 4°C for 110 days to be recovered from the VBNC state by using optimized resuscitation-buffers

| Pathogen | Microcosm | VBNC-induced period (days)<sup>2</sup> | Temperature upshift in the following resuscitation-buffers<sup>1</sup> at 25°C for consecutive 7 days |  |  |  |  |  |  |
| --- | --- | --- | --- | --- | --- | --- | --- | --- | --- |
|  |  |  | TSB | CFS-VP | CFS-VV | CFS-ST | CFS-EC | CFS-SA | CSP |
| *V. parahaemolyticus*<br>ATCC 17082 | ASW | 50 | - | 5.60 | 4.70 | 7.78 | 8.00 | 4.48 | 6.08 |
|  | ASW+5% NaCl | 25 | - | 8.75 | 5.80 | - | 5.99 | - | 6.76 |
|  | ASW+10% NaCl | 21 | 8.48 | 5.08 | 5.17 | - | - | - | 5.00 |
|  | ASW+30% NaCl | 12 | - | - | - | - | - | - | - |
| *V. parahaemolyticus*<br>ATCC 27969 | ASW | 50 | 8.08 | - | - | - | - | - | 9.15 |
|  | ASW+5% NaCl | 50 | - | 8.83 | 8.91 | - | - | - | 8.79 |
|  | ASW+10% NaCl | 50 | - | - | - | - | 8.81 | 5.94 | - |
|  | ASW+30% NaCl | 25 | - | - | - | - | - | - | - |
| *V. parahaemolyticus*<br>ATCC 33844 | ASW | 72 | 7.69 | 8.78 | 9.53 | 8.72 | 8.91 | 8.52 | 7.79 |
|  | ASW+5% NaCl | 72 | 8.91 | 7.72 | 7.46 | 8.36 | 9.00 | 7.61 | 8.26 |
|  | ASW+10% NaCl | 72 | 7.93 | 8.45 | - | 6.93 | 7.90 | 8.36 | - |
|  | ASW+30% NaCl | 25 | - | - | 8.88 | 8.28 | - | - | 6.70 |

<sup>1</sup> These pathogens in the VBNC state for 110 days were transferred into TSB (pH 8.1) supplemented with 3% NaCl, and then incubated at 25°C for 7 days.
<sup>2</sup> Days for which *V. parahaemolyticus* and *V. vulnificus* were required for the entry into the VBNC state at 4°C.

**TABLE 6.** Components of developed resuscitation-buffers

| Buffer | Component categories<sup>1</sup> |  |  |  |
| --- | --- | --- | --- | --- |
|  | TSB | CSP | MgTA | CFS |
| A | • | - | - | - |
| B | • | • | - | - |
| C | • | • | • | - |
| D | • | • | • | • |
| E | • | - | - | • |
| F | • | - | • | • |

<sup>1</sup> TSB, tryptic soy broth (pH 8) supplemented with 3% NaCl.
<sup>2</sup> CSP, 10,000 U/mg catalase and 2% sodium pyruvate.
<sup>3</sup> MgTA, 20 mM MgSO<sub>4</sub> + 5 mM EDTA.
<sup>4</sup> CFS, the cell free supernatant of *V. parahaemolyticus* ATCC 17082 grown overnight in each of appropriate resuscitation buffers at 37°C.

**TABLE 7.**
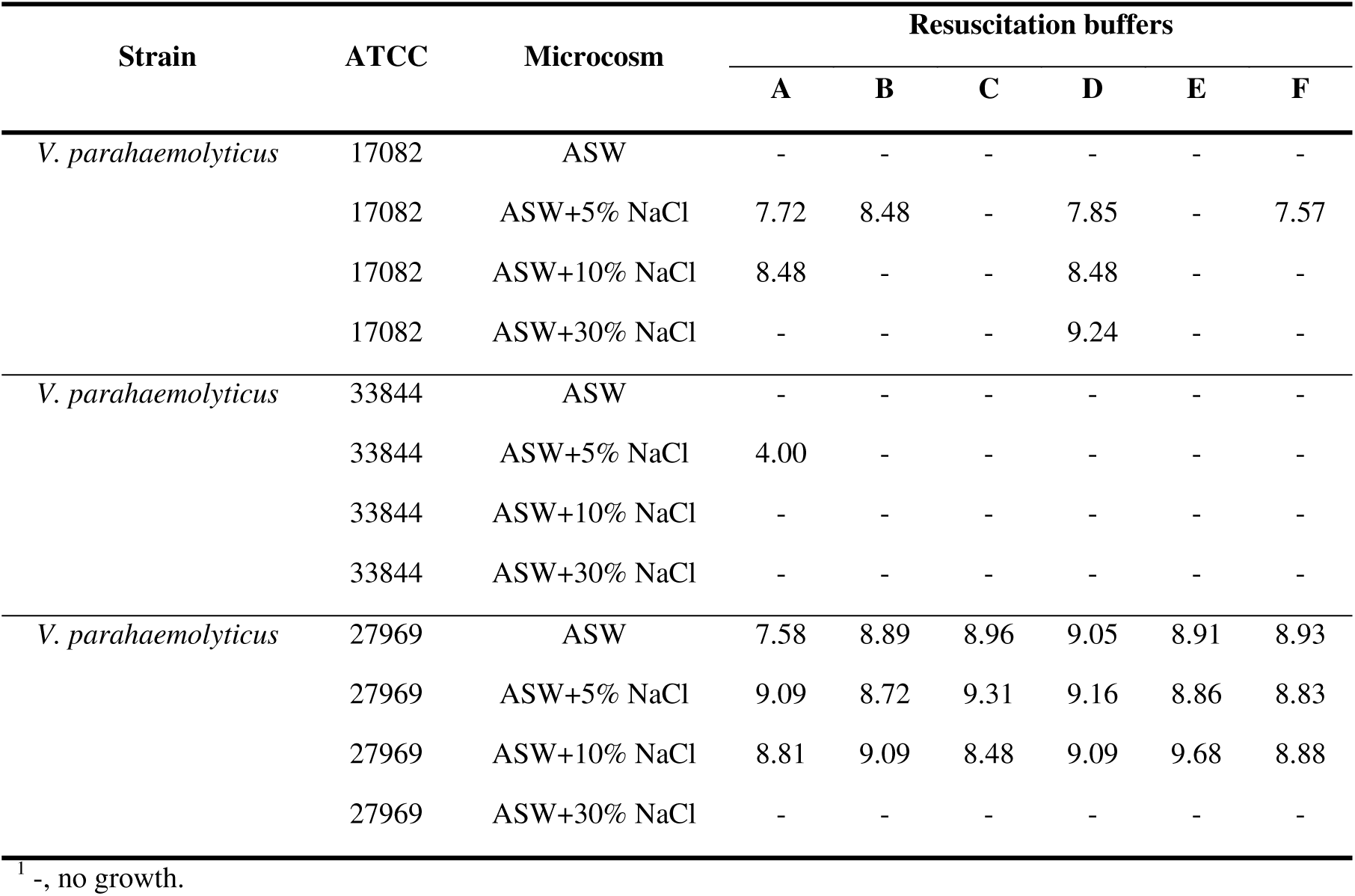
Effects of developed recovery buffers on the resuscitation of 250-days-nonculturable *V. parahaemolyticus*, followed by temperature upshift method at 25°C for 5-7 days

## ACKNOWLEDGEMENT

This research was supported by Basic Science Research Program through the National Research Foundation of Korea (NRF) funded by the Ministry of Education (Grant Number: NRF-2016R1A6A3A 11932794).

